# Generalizing about GC (hypoxia): Dys- & Dat-Informatica *Comment on* “Germinal center B cells selectively oxidize fatty acids for energy while conducting minimal glycolysis”

**DOI:** 10.1101/2021.01.28.428711

**Authors:** Mark R. Boothby, Ariel Raybuck, Sung Hoon Cho, Kristy R. Stengel, Scott Hiebert, Jingxin Li

**Affiliations:** Department of Pathology, Microbiology & Immunology, Molecular Pathogenesis Division, Vanderbilt University Medical Center and School of Medicine, Nashville TN 37232 USA; Department of Biochemistry, Vanderbilt University School of Medicine, Nashville TN 37232 USA; Medical Scientist Training Program, Perelman School of Medicine, University of Pennsylvania 3400 Civic Center Blvd, Philadelphia, PA 19104 USA

## Abstract

Steadily accumulating evidence supports the concept that the outputs of immune responses are influenced by local nutrient and metabolite conditions or concentrations, as well as by the molecular programming of intermediary metabolism within immune cells. Humoral immunity and germinal center reactions are one setting in which these factors are under active investigation. Hypoxia has been highlighted as one example of how a particular nutrient is distributed in primary and secondary follicles during an antibody response, and how its sensors could impact the qualities of antibody output after immunization. Based on a bio-informatic analysis of mRNA levels in germinal center and other B cells, recently published work challenges the concept that there is any hypoxia or that it has any influence. In this perspective, we perform new analyses of published genomics data to explore potential sources of disparity and elucidate aspects of what on the surface might seem to be conflicting conclusions. In particular, the replicability and variance among data sets derived from different naïve as well as germinal center B cells are considered. The results of the investigation highlight several broader issues that merit consideration, especially at a time of heightened focus on scientific reports in the realm of immunity and antibody responses. From one finding of this re-analysis, it is proposed that a standard should be expected in which the relationship of new data sets compared to prior “fingerprints” of cell types should be reported transparently to referees and readers. In light of the strong evidence for diversity in the constituencies within germinal centers elicited by protein immunization, it also is proposed that a core practice should be to avoid overly broad conclusions about germinal centers in general when experimental systems are subject to substantial constraints imposed by technical features.

---

In the March issue of *Nature Immunology*, Weisel, Shlomchik, and co-workers presented interesting data pioneering a use of flow-purified B cells from BCR knock-in mice to explore substrate utilization and metabolic features of classes of B lymphocytes (1). These included naïve B cells - in some but not all comparisons - and a population of germinal center (GC)-phenotype B cells recovered from recipients with a monomorphic B cell population designed to avoid inclusion of other B cells into GC (1–3). Comparisons also involved B cells after T-independent activation in vivo (1). In light of the limits to using bio-informatic data to reach conclusions about biological systems, the new evidence about fatty acid oxidation (1) advances insights beyond gene expression profiles comparing naïve and GC B cells (4). However, the paper evoked a need to evaluate the conclusive statement that the “GCBC transcriptome is not commensurate with [….] hypoxia” and similar broad conclusions of the text. This claim seems connected to a view of the authors that RNA-Seq data with GCBC do not contain evidence of enrichment for genes encoding enzymes connected to glycolysis, or that such increases relative to naïve B cells would necessarily show up in a targeted metabolomics analysis with ^13^C-labeled glucose. These issues prompted examination of these and other data sets in GEO. The results of the analyses point to limits to the conclusions as stated in (1); they also raise a broader question about the system used for this work.

As cited by the authors, several papers (4–6) have documented results from intravital labeling with imidazole compounds that covalently modify cellular constituents when the mitochondria of viable cells operate under reductive conditions due to intracellular hypoxia (7–9). Work under controlled conditions has shown that a meaningful signal above background is obtained only when the ambient pO_2_ is below a level sufficient to yield HIF stabilization (7–9). Direct evidence of increased HIF-1α has been presented for both GC B cells (4, 5) and their Tfh counterparts (10). Any issue of HIF function requires understanding that TCR and BCR engagement cause sustained HIF-1α and HIF-2α stabilization, which presents a drawback to comparisons between activated and GC B cells. It might formally be possible that duration of the hypoxia, BCR signaling, and HIF stabilization failed to yield changes in mRNA concentrations large enough for enough gene products to yield a “statistically significant” result in a gene set enrichment analysis (GSEA) algorithm. Indeed, using a gene signature derived with the human breast cancer-like cell line MCF7 (11), the authors’ analysis of their purified GC B cells suggested that neither hypoxia-related nor HIF-1 target genes were enriched under their conditions of experimentation. Moreover, application of a different gene signature indicative of biologically significant hypoxia (12) to their data sets also yielded a balanced mix of increased versus decreased mRNA, in contrast to a previously published analysis in which GSEA scored the result as statistically significant (4). This difference along with other disparities of the data prompted us to compare the informatics findings while also using additional benchmarks that could test each report for independent replication.

Hypoxia signal was observed with several types of immunization, including with NP-carrier (ovalbumin) (6), but work apart from (4) did not have RNA-Seq data for comparisons. However, contemporaneous (2016, 2017) papers with data deposited in GEO have replicate data on naïve and bulk GC B cells purified in a similar time frame (7-10 d) after immunization with the same immunogen (SRBC) (13, 14). The sequencer output data for all four papers (1, 4, 13, 14) were analyzed using the same pipelines (one assembled in-house, detailed in a technical log appended to this note, and a second via Basepair Technologies, Inc for an independent framework). Several salient observations emerged from these comparisons. Unsupervised clustering with Spearman correlation analyses and the Euclidean distances among the different types of samples (**Fig. 1**) show the results from (1) to differ substantially from the data of (3), (13), and (14). *First*, the mRNA expression pattern of GC B cells generated after transferring large numbers of B cells [apparently, 10^6^ - (2) as cited in (1)] biased toward a single specificity BCR (B1-8^i^ Vκ−/−) into recipient mice with a monomorphic B cell population specific for Ig (AM14-Tg × Vκ8R-KI BALB/c mice, i.e., specific for allotype-disparate IgG2a^b^) differed substantially from the other samples. A measure of overall differences was less between naïve and GC B cells in (1) (mean Euclidean distances of 126.3 ± 0.69) than the difference between the GC B cells of the transferred B1-8i, Vκ−/− cells (1) and the polyclonal GCBC (4, 13, 14) (mean Euclidean distances of 170.3 ±10.3; p<0.01) as well as those of non-transgenic naïve versus GC B cells (147.6 ± SEM of 3.3; p<0.05). *Second*, the GC B cells from mice with a normal pre-immune repertoire (4, 13, 14) were substantially more similar to one another in comparing among three independent analyses (correlation coefficients in the range of +0.41 to +0.92) as opposed to anti-correlation of the B1-8i-derived GCBC, (correlation coefficients of −0.05 to −0.32). In addition, the RNA-Seq “fingerprints” of naïve B cells in a polyclonal system were substantially similar to each other. Thus, the data were replicable when comparing independent SRBC immunizations of mice with BCR or limitation of the repertoire. In contrast, even the naïve B1-8^i^ Vκ−/− B cells were quite distinct from the clusters of polyclonal naïve B cells. Relative to the data from (14) as well as (4), the changes (Euclidean distances) were more modest in (1). To some extent, the similarities and differences among independent data sets can be parsed by heat maps of differentially expressed genes among the data sets (**Fig. 2A**) and visualization of Principal Components Analyses (**Fig. 2B**), albeit with the caveat that the simplification (dimensionality reduction) of principal components reduces detail. In practice, the collective data suggest that the GCBC generated after adoptive transfers of B cells with a restricted repertoire into mice whose GC will bring in limited endogenous B cells are qualitatively distinct from a polyclonal response that evolved from a polyclonal repertoire. Several factors may be involved in the difference highlighted by these data. Akin to findings with T cells (15), they may include the intra-clonal competition that can result from using hundreds of thousands of transferred antigen receptor-transgenic cells (16) - as already documented for B1-8i (17). There also is the potential for highly diverse GC to have differences in their metabolic environment from the more uniform secondary follicles that would ensue with the transfer system used in (1). These factors warrant investigation, and it should not be misunderstood that these analyses of differences are to say that one answer using a very different approach is “wrong” when conclusions are restricted to that approach. Instead, the key points indicated by the data are that the cells analyzed in (1) changed less from naïve to GC phenotype than what is found in reproducible data with B cells that derived from a normal polyclonal repertoire (1, 13, 14). In part, these B1-8i, Vκ−/− B cells started out with substantial differences (including in an activation marker, *Nr4a1*, and differentiation-related genes, i.e., *Prdm1* and *Irf4*).

**Figure 1.**
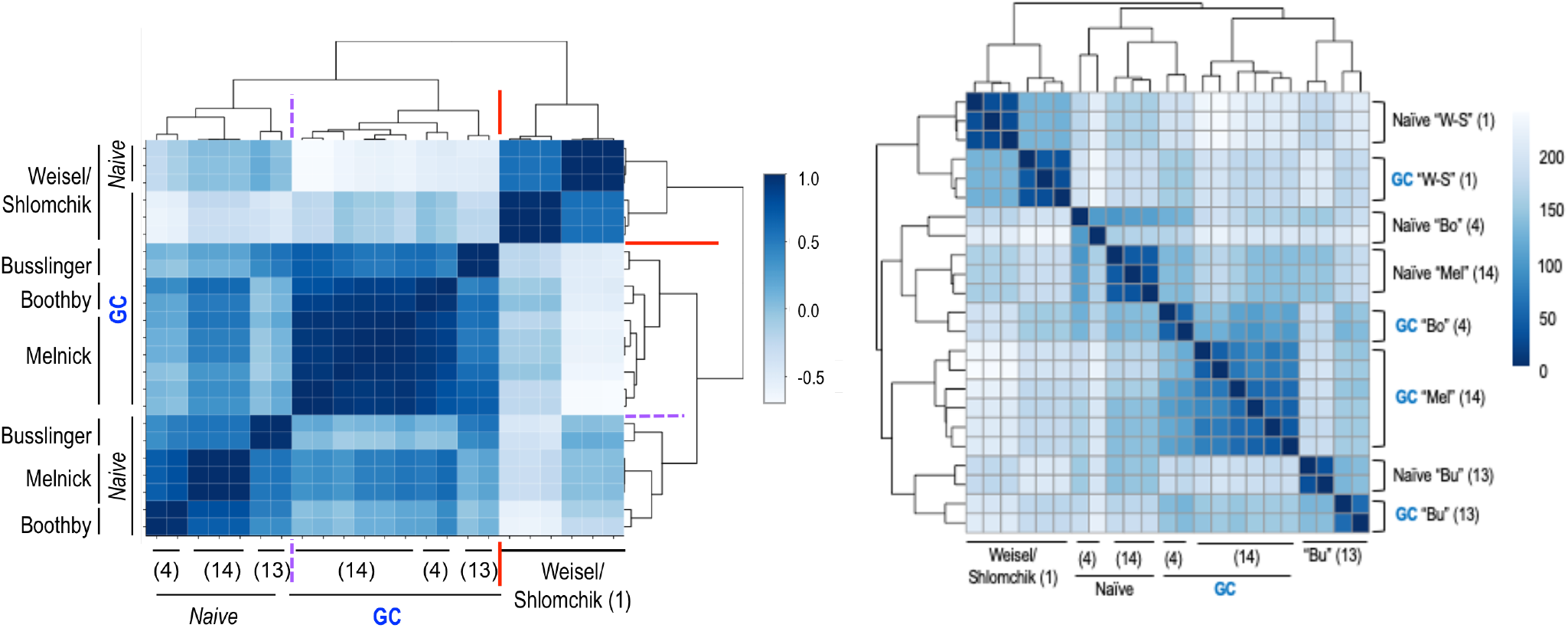
Spearman correlation and Euclidean distance analyses applied across the RNA-Seq datasets of naïve and GC B cells in the transgenic and non-transgenic systems. Raw RNA-Seq data for naïve and GC B cells were downloaded from the GEO deposits for the papers cited as (1), “W-S”; (13), “Bu”; (4), “Bo”; (14) “Mel”, and put through each of two analysis pipelines applied uniformly to all data (detailed in the Methods and technical log). (*Left*) Self-organizing map from unsupervised clustering based on Spearman correlation coefficients across the data sets of the papers cited as (1), (4), (13), and (14). Darker blue represents strong positive correlation (1.0 = identical across the RNA-Seq data); lightest blues are anti-correlated. (*Right*) Self-organizing map from the Euclidean distances between a particular sample and comparator samples. Darkest blue, at “0”, denotes completely identical values across the RNA-Seq data (i.e., a sample compared to itself). Higher numbers denote lower similarity (greater difference, i.e., greater Euclidean distance, across the gene expression “fingerprint”).

**Figure 2.**
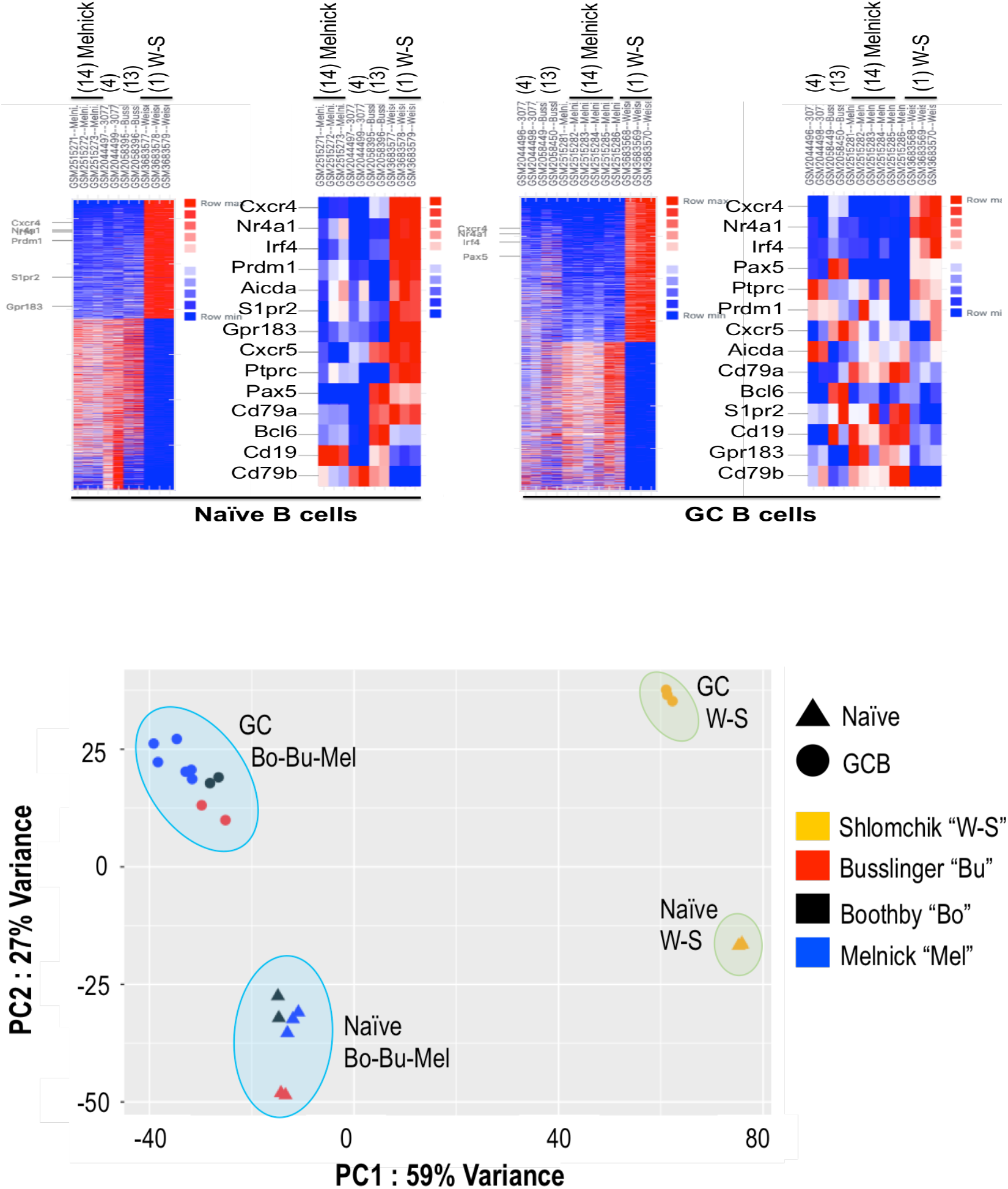
A. Comparisons of condition-dependent differential gene expression. B. Principal components analysis of the RNA-Seq datasets of naïve and GC B cells in non-transgenic systems versus B1-8i, Vk −/− transgenic cells into AM14. GEO data deposited for the papers cited as (1), “W-S”; (13), “Bu”; (4), “Bo”; (14) “Mel” were used for (A) differential expression and (B) Principal Component Analyses (PCA). (A) Heat maps are shown for naïve and GC B cells, with either all differentially expressed genes (16% and 14% of totals, respectively, i.e., 3214/22,843 and 3461/21,327 genes differentially expressed >2-fold with p-adjusted <0.05) or a subset of B lineage-specific or other genes functionally relevant in GC biology. (B) PCA plot depicting results across the mined datasets for the indicated conditions (shape) and datasets (color). X- and Y-axes are defined by PC1 and PC2, accounting for 59% and 27% of the variance across the datasets, respectively. Further details are in a Methods and technical log (below).

As to an underlying question, are there hypoxia-related gene signatures in GC B cells when taking into account other work with a polyclonal repertoire? The application of GSEA algorithms when using several different gene sets for hypoxia yields statistically significant results (**Fig. 3**). Moreover, RNA-Seq data with GC from NP-CGG-immunized mice with a normal pre-immune landscape (20) also show enrichment for a functional hypoxia signature (**Fig. 3**, data from the Pernis lab), and divergence from the B1-8i data. GC are quite heterogeneous (18, 19). This point suggests that caution may be warranted from the outset, militating against conclusions being framed as blanket generalizations to apply across all GC. In terms of hypoxia or HIF gene signatures, prior work reported (4) that as many as 20% of splenic GC had no signal indicative of intravital hypoxia, a variegation observed in parallel by others (J. Jellusova, personal communication). An integrative possibility is that hypoxia and/or its influence are reduced in GC dominated by a single specificity and designed to minimize bystander B cell involvement as in (1–3). Two further facets of what the analyses reveal are that *(i)* with some hypoxia modules, even the Weisel-Shlomchik data show significant enrichment in GSEA, and *(ii)* there is not really a validated hypoxia or HIF module for lymphocytes, let alone a unitary module that includes activated or GC B cells. Although hypoxia and HIF responses are protean, if three decades of papers on cell type-specific functionality of a transcription factor apply to GC or B cells, some restraint may be warranted before concluding that a GC transcriptome is not commensurate with hypoxia. The same caution might apply to interpreting the changes in mRNA levels for glycolysis-related genes. Prior work had provided evidence that the “glycolysis” and “FAO” transcriptomes were increased in GCBC when compared to naïve B cells (4). Although this particular comparison was not evident in the Figure presented in (1) using the B1-8i system, a GSEA on naïve and GCBC with the RNA-Seq data in (1) shows significant increases in glycolysis-related gene expression, which also is observed in each of the other data sets. Of course informatics analyses – conflicting or not – cannot settle the issue of glucose uptake and utilization. Translation efficiency, post-translational modifications, and complex but unknown aspects of nutrient supplies, substrate concentrations, and substrate competition along webs of interconnected pathways all are downstream from the mRNA in question. That simply means that direct rigorous biochemical assays of glucose uptake per cell [not just 2-NBDG, which does not necessarily correlate with glucose entry rates (21)] and glucose oxidation are needed before reaching broad conclusions about germinal center B cells based on a single or special case. In this respect we suggest that normalizing to cell size falls beyond the pale of rigor - unless it would be accepted that a large truck does not consume more fuel than a compact sedan because the size-normalized fuel consumption is the same, or that a 160 kg person eating 4400 kCal/d is taking in the same amount of energy as one who is 80 kg and eats 2200 kCal/d. In regards to a conclusion based solely on informatics, it is proposed that a reasonable standard is to perform a comparison of new RNA-Seq data sets to independent entries in GEO, and to quantify the correlation to - or difference from - those that are in GEO and readily comparable (i.e., not micro-array to RNA-Seq). Importantly, some restraint in statement of conclusions and openness about limitations seems particularly vital at a time when the societal landscape is roiled by concerns about how scientists frame their work or evidence.

**Figure 3.**
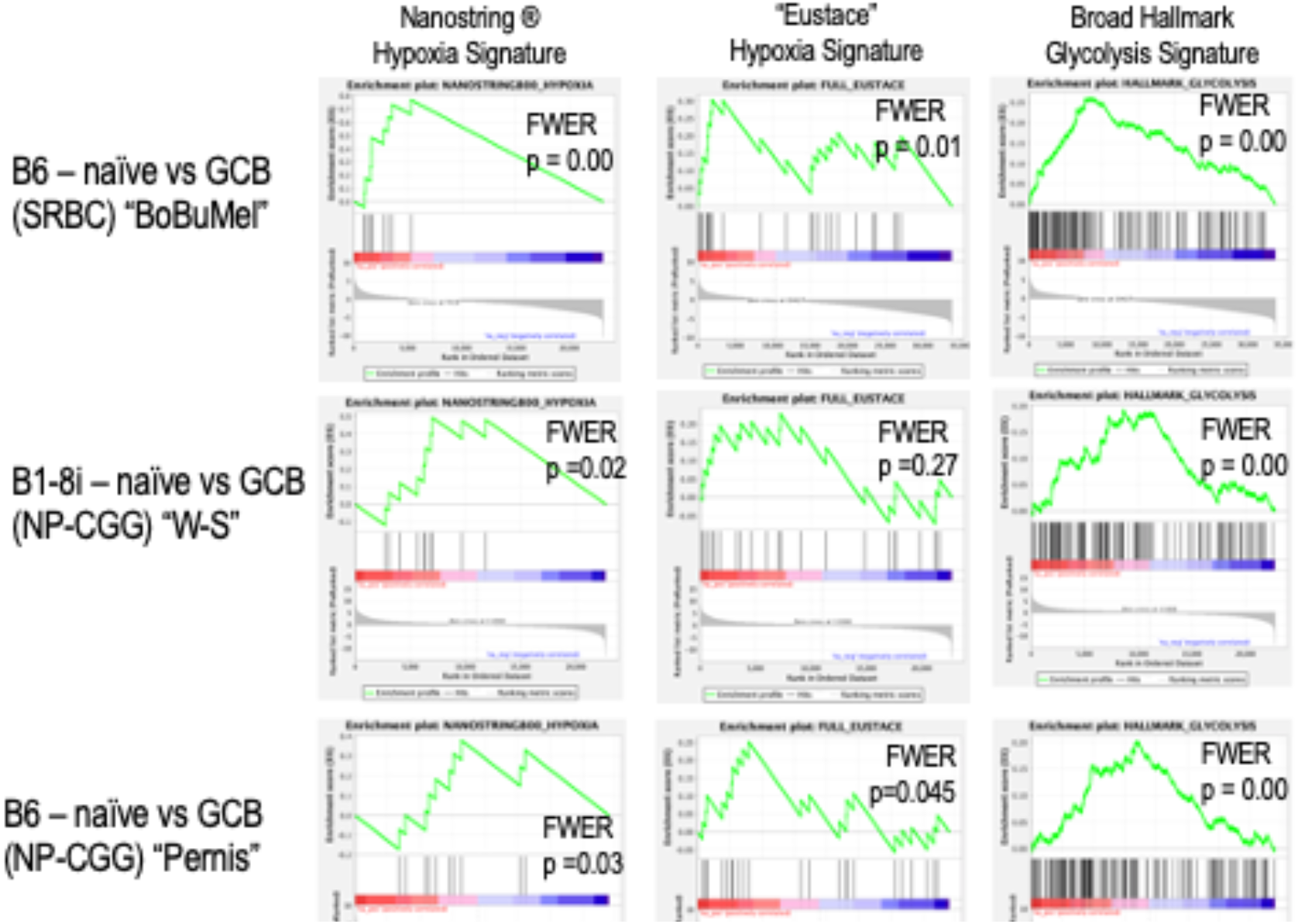
GSEA with the RNA-Seq datasets of naïve and GC B cells in the transgenic transfer model versus a meta-analysis of the non-transgenic systems. Raw data from RNA-Seq analyses of the cited papers (references 1, 4, 13, 14, and 20, using name or initials of the senior authors as in Fig. 1, 2) were processed through a pipeline analogous to that of (1). Each panel displays results for a separate analysis in which ranked differential expression data were processed using the Broad Institute’s GSEA software. Insets in each panel display the p values derived after corrections for multiple comparisons (FWER). The three columns represent outputs obtained using the indicated gene signatures: Hypoxia Signature as established by Nanostring Technologies ®, the hypoxia gene signature of Eustace et al (as in references 4, 12), and the Broad Institute’s Hallmark Glycolysis geneset. The rows of panels identify the RNA-Seq data sets that were analyzed in comparisons of GC to naïve B cells: (i) data from like samples in (4), (13), and (14); (ii) data from (1); and (iii) data from (20), all of which were processed through the same pipeline with same parameters. Steps in the analyses are detailed in the Methods log.

## Methods and technical log

Datasets comparing RNA expression data from “naïve” (IgD+ or Follicular) versus immunization-induced GC B cells were mined from the GEO depositions of raw sequencing data that all were generated with Illumina Hi-Seq 2000 or 2500 instruments. Sequences were trimmed using the fastp FASTQ preprocessor for overrepresented sequences and to remove any sequence with a quality score <10 (1). Trimmed sequences were aligned to the mm10 mouse genome using the STAR sequence aligner of Dobin et al (2). Aligned sequences were then quality-tested using Qualimap (3) software, and counted using featurecounts in Rsubread (4). There were no substantive differences between different data sets in the quality [for naïve and GC B cell RNA respectively, 92% and 91-92%, 95% and 94-95%, 94% and 93%, and 87% and 91% for the papers referenced in the main text as (1, 4, 13, and 14, respectively) via the in-house pipeline uniformly applied to all samples. Comparable values, all above 90%, were generated by the Basepair Technologies pipeline and STAR alignment algorithms. Length scoring also was indistinguishable among the different datasets, two of which [the papers cited as (1) and (14) in the main text] were generated by paired-end sequencing. These steps were followed by multi-factor differential expression analysis in which deposited datasets as well as “naïve” vs “GCB” expression data were compared using DESeq2 (5). Effect size shrinkage of differential expression data was normalized using the *apeglm* method published by Zhu et al (6). Processing of the gene expression and PCA was cross-checked by using the commercial pipeline of Basepair Technologies as applied to the primary data for naïve and GC B cells of the papers cited as (1, 4, 13, 14). Gene set enrichment analyses were carried out using the GSEA program of the Broad Institute (7,8), and gene sets derived from published literature, the Rat Genome database (“RGD”), and the Broad KEGG database. Euclidean distances and Spearman correlation coefficients were derived by applying a standard “dist” function in R to the data described above.

